# Characterization of intact proviruses in blood and lymph node from HIV-infected individuals undergoing analytical treatment interruption

**DOI:** 10.1101/456020

**Authors:** Line K. Vibholm, Julio C.C. Lorenzi, Joy A. Pai, Yehuda Z. Cohen, Thiago Y. Oliveira, John P. Barton, Marco Garcia Noceda, Ching-Lan Lu, Yuria Ablanedo-Terrazas, Perla M. Del Rio Estrada, Gustavo Reyes Teran, Martin Tolstrup, Paul W. Denton, Tine Damsgaard, Ole S. Søgaard, Michel C. Nussenzweig

## Abstract

The role of lymphoid tissue as a potential source of HIV-1 rebound following interruption of antiretroviral therapy is uncertain. To address this issue, we compared the latent viruses obtained from CD4^+^ T cells in peripheral blood and lymph nodes to viruses emerging during treatment interruption. Latent viruses were characterized by sequencing near full-length (NFL) proviral DNA, and *env* from viral outgrowth cultures (VOAs). 5 HIV-1 infected individuals on antiretroviral therapy (ART) were studied, 4 of whom participated in a clinical trial that included an analytical treatment interruption. Intact or replication competent clonal sequences from blood and lymph node overlapped. In contrast, there was no overlap between 205 latent reservoir and 125 rebound sequences in the 4 individuals who underwent treatment interruption. However, rebound viruses could be accounted for by recombination. The data suggests that CD4^+^ T cells carrying latent viruses circulate between blood and lymphoid tissues in individuals on ART and support the idea that recombination may play a role in the emergence of rebound viremia.

## Introduction

ART is an effective treatment for HIV-1 infection. However, HIV-1 persists as proviral DNA in CD4^+^ T cells and produces a latent reservoir that demonstrates remarkable stability (1–3). Upon ART interruption, nearly all individuals experience viral rebound within 1-6 weeks (4–6). However, most integrated proviruses are defective (7, 8), and only a small percentage can replicate and produce the infectious virions that mediate rebound viremia when ART is interrupted. Latent viruses can be recovered from the reservoir by viral outgrowth assays (VOA) and by polymerase chain reaction amplification of integrated proviruses (9, 10). Using these methods latent viruses have been recovered from lymph nodes, gut-associated lymphoid tissue (GALT), spleen, CNS, liver, lungs, kidney, adipose tissue and genital tract(11–17).

In 4 recent studies, comparison of circulating latent viruses and rebound viruses showed a very limited number of overlapping sequences (10, 18–20). One potential explanation for this observation is that latent viruses found in CD4^+^ T cells in lymphoid and other tissues differ from those in circulation, and that these sequestered cells are responsible for the rebound viremia. To examine the relationship between latent viruses in circulation and in lymph node, we studied 5 individuals who had concurrent blood draws and lymph node biopsies. Rebound plasma was available for 4 of these individuals who were enrolled in a clinical trial that included an analytical treatment interruption after 24 weeks of therapy with a TLR9 agonist.

## Results

### Samples and study participants

To investigate the intact proviral reservoir, we obtained mononuclear cells from lymph nodes (LNMC) and peripheral blood (PBMC) from 4 ART treated HIV-1 infected individuals (extended data table 1a), who had been virally suppressed for a median of 8.3 years (range 1.8–13.4). All 4 participated in an interventional trial in which a TLR9 agonist was co-administered with ART for 24 weeks (figure 1a) (21). PBMCs and LNMCs were collected in the last 2 weeks of the 24 week period. Subsequently ART was interrupted until viral rebound. Plasma was collected before ART re-initiation. Time from ART withdrawal to viral rebound was between 9–15 days (figure 2a) which is not significantly different from a non-interventional ATI control cohort of 52 participants (ACTG cohort) (22–24) (figure 2b, P = 0.5, Log-rank test). In addition, we obtained PBMCs and LNMC from an HIV-1 infected individual (ID LFSO), who had been on ART for 1.3 years without additional interventions or treatment interruption (extended data table 1b). To obtain full-length *env* sequences from viruses that were replication competent and/or genetically intact, CD4^+^ T cells from blood and lymph node were assayed by VOA, and near full-length (NFL) PCR (10) (extended data table 2).

**Figure 1.**
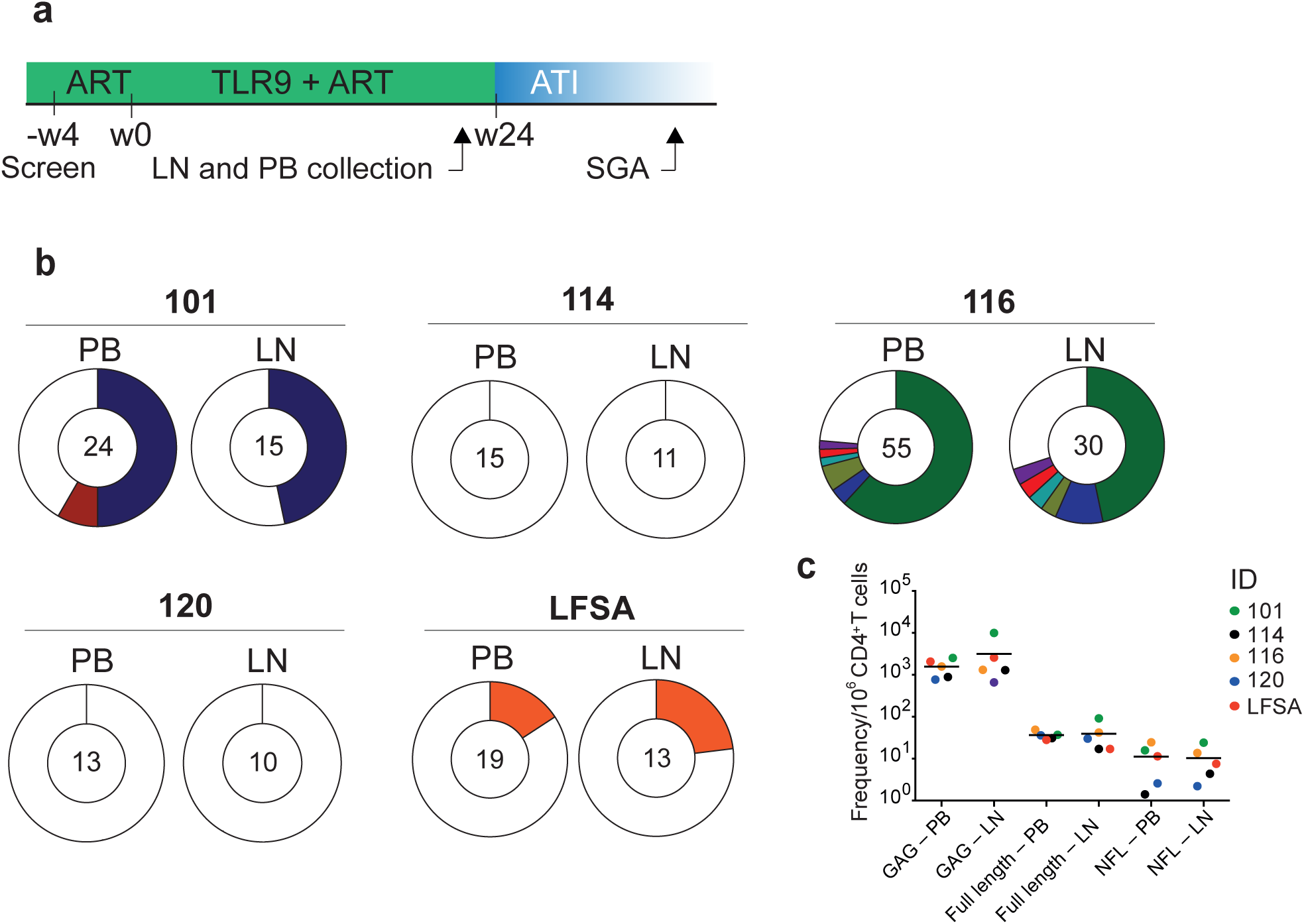
TLR9 agonist study design and clonal distribution of the latent HIV-1 in peripheral blood and lymph nodes. **a**, Study design. Green area represents time on ART before enrollment and during the 24 weeks of TLR9 agonist treatment. Blue area represents time off ART. “W” and numbers and represents weeks elapsed. Lymph node (LN) and peripheral blood (PB) were collected 1-2 weeks before initiation of the analytical treatment interruption (ATI). SGA depicts the time point where plasma was collected and ART subsequently reinitiated. **b**, Pie charts showing the distribution of intact near full-length (NFL) and VOA-derived *env* sequences. Numbers in the center of the circles represents total number of intact/replication competent sequences obtained. White areas in pie charts are sequences obtained once (singles). Colored areas represent sequences obtained more than once (clones). Clones, which are found in both PB and LN within an individual, share the same color between the two PB/LN pie charts. The size of the slices in the pie charts is proportional to the relative size of the clone. For ID 101, 116 and LFSA, there is no significant difference between the frequency of clones in PB and LN (two-sided Fisher’s exact test). **c**, Frequency of GAG+ cells per 10^6^ CD4^+^ T cells in LN and PB. Frequency of full length viruses (amplicon size determined using 0.8% agarose gel: i.e. at least one combination of either A+C, A+D, B+C or B+D possible (Ho *et al.*, 2013)) per 10^6^ CD4^+^ T cells in LN and PB. Frequency of intact near full-length (NFL) sequences per 10^6^ CD4^+^ T cells in LN and PB. There was no statistical difference between the frequencies of GAG+, (p-value=0.31) full-length viruses (p-value=0.63) and intact NFL (p-value=0.81) per 10^6^ CD4^+^ T cells (Wilcoxon matched-pairs signed-rank test). Each participant has a unique color code.

**Figure 2.**
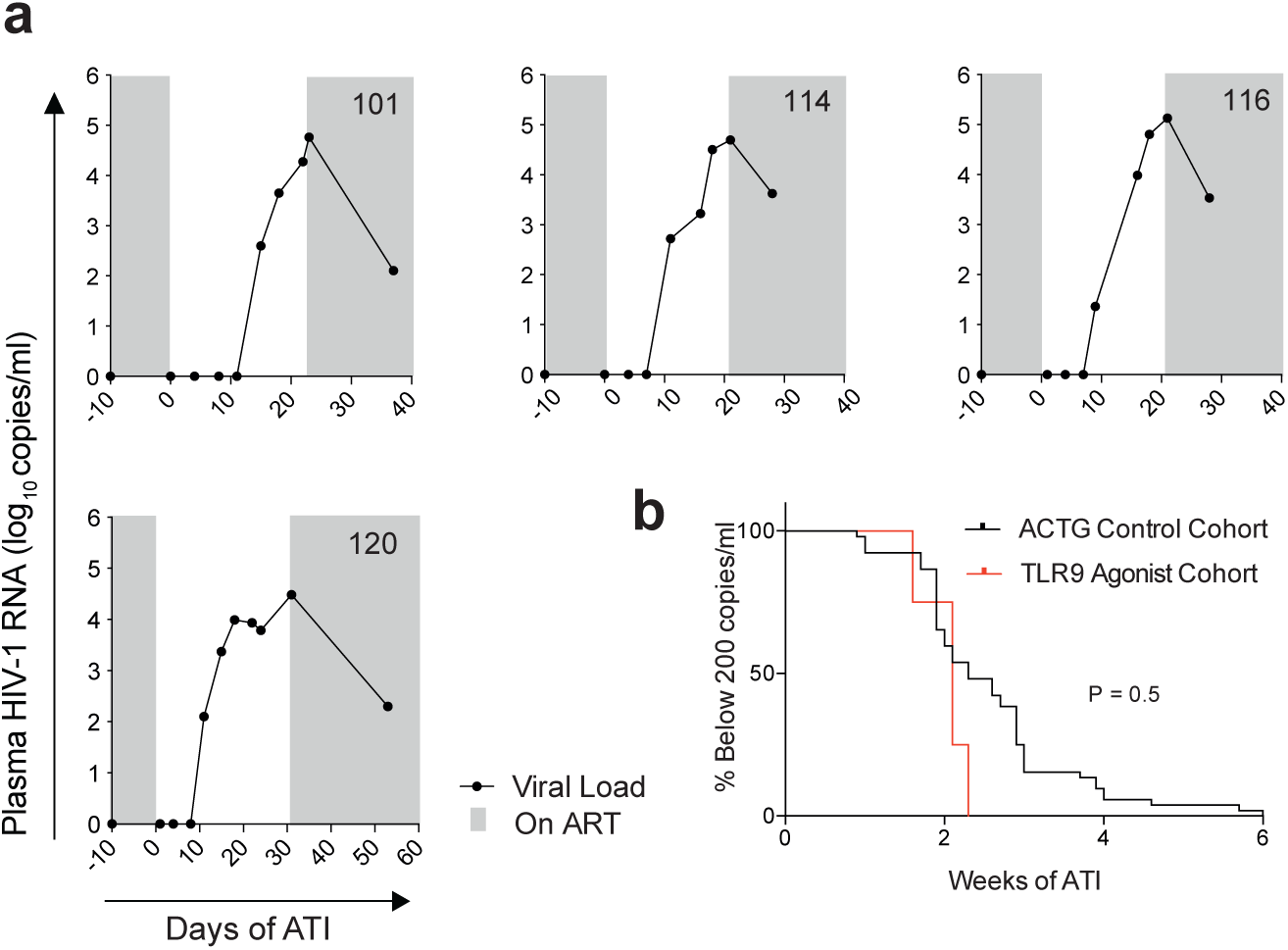
Time to viral rebound during ATI. **a,** Plasma HIV-1 RNA levels (left y-axis) and days elapsed (x-axis) for the 4 participants receiving the TLR9 agonist. Black lines on the x-axis connect dots that denote HIV-1 RNA levels at the indicated number of days after ATI. Lower limit of HIV-1 RNA detection was 20 copies/mL. Gray shaded areas depict time on ART. **b**, Kaplan-Meier plot summarizing time to rebound for the 4 TLR9 agonist trial participants (red line) compared to a cohort of 52 ACTG trial participants (black line) who underwent ATI without intervention. Logrank test P value shows the comparison of time to rebound for the TLR9 agonist treated and the ACTG cohort.

### Intact and defective latent viruses from blood and lymph node CD4^+^ T cells overlap

We obtained 205 latent virus sequences: 93 and 79 intact NFL sequences from blood and lymph node, respectively, and 33 from VOA from blood (extended data table 2). 44% of all sequences belonged to expanded clones, 98% of which overlapped between blood and lymph node (i.e. identical sequences). There were no statistical differences between the frequency of clonal sequences in blood compared to lymph node (two sided Fisher’s exact test, p-values in extended data table 3a). Clones were absent in participants ID 114 and ID 120 where sample availability was more limited and latent viruses were diverse.

To determine whether defective viruses were also similar between blood and lymph node, we combined intact and defective *env* sequences from all NFL and VOAs (327 sequences) (extended data figure 1). Overall 43% of all *env* sequences belonged to expanded clones (143 sequences), and 89% of the clones overlapped between blood and lymph node (two sided Fisher’s exact test, p-values in extended data table 3b). We conclude that CD4^+^ T cells in peripheral blood and lymph node contain overlapping sets of proviruses.

### The frequency of intact viruses in blood and lymph node

To further examine the relative proviral nucleic acid content in peripheral blood and lymph node, we compared the frequency of: 1) GAG+ cells; 2) full-length genomes; and 3) intact NFL proviruses (figure 1c). The frequency of GAG+, full-length genomes and intact NFL per 10^6^ CD4^+^ T cells was not statistically different between peripheral blood and lymph node (figure 1c). Thus, in our cohort the frequency of intact and defective HIV-1 proviruses in CD4^+^ T cells is similar in peripheral blood and lymph node.

### Relationship between plasma rebound viruses and reservoir viruses

125 full-length *env* sequences were obtained by SGA from rebound plasma, none of which overlapped with any of the latent reservoir sequences (figure 3). Nevertheless, phylogenetic analysis revealed that plasma rebound viruses were related to latent viruses in peripheral blood and lymph node from the same individual (figure 4a and 4b). Thus, the latent viruses obtained from peripheral blood and lymph node are related to, but do not appear to be the direct origin of rebound virus in these individuals.

**Figure 3.**
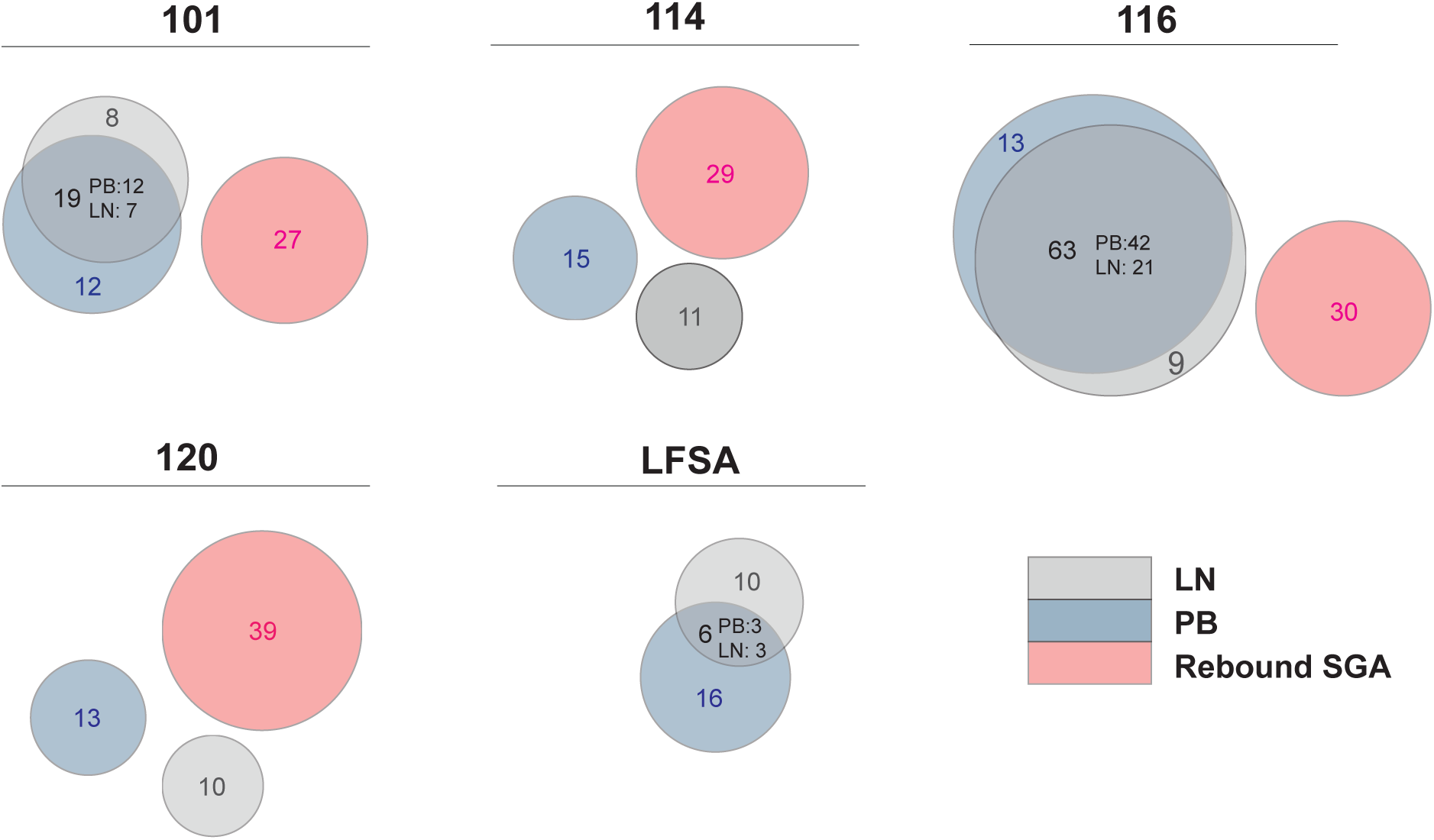
Sequence identity between peripheral blood, lymph node and rebound single genome assay (SGA) viruses. Venn diagrams depicting env sequences from peripheral blood (PB) near full-length (NFL) and PB VOA (blue), lymph node (LN) NFL (gray) and SGA from the time of viral rebound (pink). Number of sequences obtained are indicated in the circles. The relative size of the overlapping areas is proportional to the number of identical sequences.

**Figure 4.**
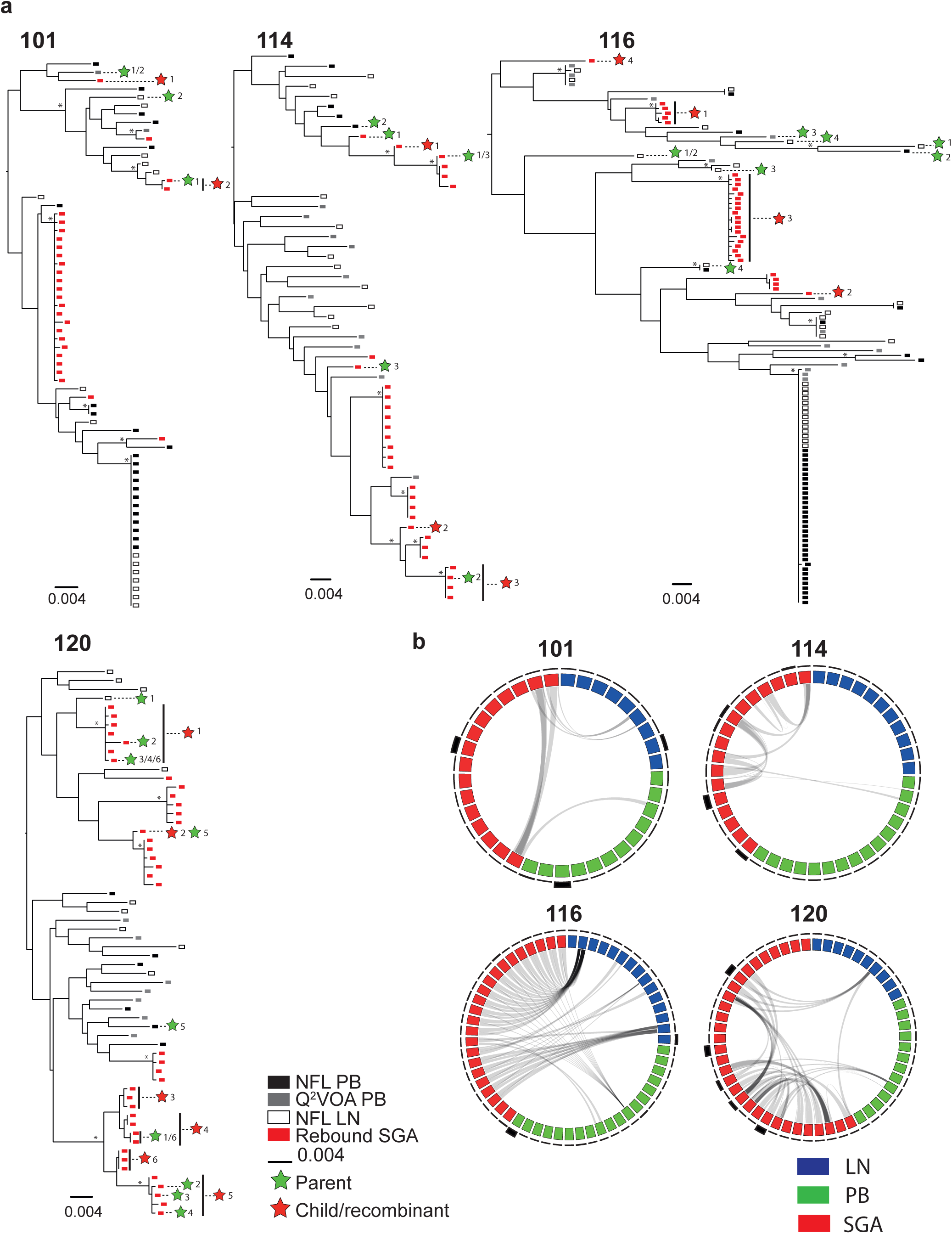
Comparison of *env* from intact sequences obtained from lymph node, peripheral blood cells and rebound viruses. **a**, Maximum likelihood phylogenetic trees of *env* from near full-length (NFL) lymph node (LN) and peripheral blood (PB) sequences, viral outgrowth (VOA) PB culture sequences and plasma SGA sequences. Symbols are defined in the graph legend. Asterisks indicate nodes with significant bootstrap values (bootstrap support ≥ 90%). Green stars are parent sequences, which undergo recombination to produce a child sequence (red star). Each recombination event has a number (denoted next to the colored star). **b,** Circos plots showing the connection between the two parent sequences and the child/recombinant sequence which are also depicted in the trees. The blue blocks represent latent reservoir LN sequences. Green blocks represent latent reservoir PB sequences. Red blocks are plasma virus sequences. Clonal sequences are depicted once. The density of the outer black lines surrounding the circles represents number of sequences retained within the clone, i.e. thin black lines are singles and thicker lines represents sequences obtained several times. Gray lines inside the circos plots shows the recombination event. Parent/child relationship is shown in the trees.

To determine whether mutations accumulating during rebound can account for the divergence between the latent reservoir and rebound viruses we used a stochastic mutation simulation model (20, 25, 26) (extended data figure 2). We found no instance in which rebound sequences could be accounted for by mutation. To determine whether recombination between latent blood and lymph node viruses could account for the rebound viruses we analyzed intact NFL, and VOA *env* sequences using the 3SEQ recombination algorithm (http://mol.ax/software/3seq/). Rebound viruses in all 4 individuals showed evidence of recombination (figure 4a, b, i.e. rejection of the null hypothesis of clonal evolution; p-values in extended data table 4 and 5). By including the possibility of mutation and recombination, we were able to account for 53% of the rebound sequences.

## Discussion

HIV-1 proviruses are present in lymphoid cells in all tissues analyzed to date (11–17). However, the relationship between viruses found in different sites and their contribution to persistence is not well defined. Our data suggest that the latent viruses found in circulation overlap with those found in lymph nodes and that the most prevalent viruses in these compartments are not typically found among rebound viruses during treatment interruption.

It has been suggested that lymph nodes serve as a sanctuary site for latent HIV-1 (27, 28). However, we found the same overall frequency of HIV-1 proviruses in blood and lymph node CD4^+^ T cells. Moreover, the two compartments contained similar numbers of replication competent viruses, and shared clones of latent viruses. Thus, expanded CD4^+^ T cell clones bearing latent viruses circulate between these two compartments. Finally, rebound viruses did not overlap with either blood or lymph node, but instead appeared to represent recombinants.

Our findings are consistent with a number of previous observations indicating that there are similar amounts of HIV-1 DNA in CD4^+^ T cells in blood and in tissues (29, 30). However, our work is limited by the numbers of individuals and CD4^+^ T cells analyzed and therefore we cannot rule out the possibility that a subset of lymph node cells, such as CD4^+^ TFH cells, are enriched in or harbor a specific group of latent viruses (31).

Three other studies have also shown little or no overlap between circulating latent viruses and rebound viruses (10, 19, 20). Including the data reported here, there are only 3 overlapping sequences among 1816 independently derived latent reservoir viruses and 642 rebound viruses. Instead, the latent and rebound compartments appear to be related by recombination.

There are a number of potential explanations for the lack of concordance between latent and rebound viruses. One possibility is that latent reservoir sampling has been inadequate. For example, the active reservoir responsible for rebound might be found primarily in a tissue that has not been assayed, such as the gut. A second non-exclusive possibility is that the majority of the latent viruses in blood and lymph node fail to emerge *in vivo* because they are in some way unfit to do so. For example, the majority of latent viruses assayed could be susceptible to immune pressure resulting in selection of a subset of rebound viruses which can escape anti-viral immunity *in vivo* possibly by recombination (32, 33).

In conclusion, the data reported indicated that the majority the latent clonal viruses found in blood are also found in the lymph nodes and add to the growing body of literature suggesting that rebound viruses are either not present in or are rare components of the latent reservoir found in circulation.

## Materials and Methods

### Study Design and participants

This study was approved by the Danish Research Health Ethics Committee (1-10-72-133-17) and the Danish Data Protection Agency, by The Rockefeller University Institutional Review Board (protocol TSC-0910) and Research Committee and Research Ethics Committee of the Instituto Nacional de Enfermedades Respiratorias "Ismael Cosío Villegas". Samples for this study were generated from a clinical trial conducted in 2016–2017 at Aarhus University Hospital, Denmark (21) (clinicalTrials.gov identifier: NCT02443935). The clinical study was approved by the Danish Research Health Ethics Committee (case no: 1-10-72-10-15), the Danish Medicines Agency (2015014125) and the Danish Data Protection Agency. Participants were recruited from the Department of Infectious Diseases Outpatient Clinic at Aarhus University Hospital and signed a written informed consent before any study procedures. Inclusion criteria were plasma HIV-1 RNA <50 c/mL, CD4^+^ T cell count >350 cells/µl, age >18 years, on cART for >12 months and ability to provide informed consent (a complete list of inclusion and exclusion criteria is accessible at clinicaltrials.gov). Inclusion criteria for participation in the ATI were: HIV RNA< 20 c/mL, CD4^+^ T cells > 350 cells/µl and written informed consent to withdrawal of cART. COBAS TaqMan HIV-1 Test, (version 2.0 by Roche) was used to assess plasma viral load (pVL) x2/week and CD4^+^ T cell count was measured every other week. Participants were called in for re-initiation of ART and collection of plasma after 2 measurements of HIV-1 RNA >5000c/mL or CD4^+^ T cells < 350 cells/µl. Data on the historical control cohort (ACTG trial: ACTG 371(22) A5024(23), A5068 (24), and A5197(22)) were used to compare to viral rebound data from the 4 participants in this study. The studies were carried out as an ATI without any further interventions and included 52 participants. Inclusion criteria for the ACTG cohort were: age 18–65, on cART>12 months, plasma HIV-1 RNA<50 c/mL>12 months before ATI initiation, CD4^+^ T cell count at time of ATI initiation>500 cells/µl, nadir CD4^+^ T cell count>200 cells/µl. Viral load was measured weekly until viral rebound occurred.

### Viral outgrowth assay (VOA)

The VOA was performed using PBMCs from peripheral blood (PB) as previously described with some modifications (34). Briefly, PBMCs were isolated by density centrifugation on Ficoll (Thermo Scientific). CD4^+^ T cells were isolated from the cryopreserved PBMCs through negative selection with magnetic beads (Miltenyi). Purified CD4^+^ T cells (0.1 ×10^6^) were cultured together with 0.2×10^6^ irradiated heterologous PBMCs from an HIV-1 negative donor in 200ul media [RPMI 1640 (Gibco) supplemented with 10% FBS (HyClone; Thermo Scientific)], 1% penicillin/streptomycin (Gibco), 1 μg/mL phytohemagglutinin (Life Technologies), and 100 U/mL IL-2 (Peprotech)] at 37°C and 5% CO2. At this density less than 30% of cultures became p24-positive. Cultures rested overnight and after 24 h, 125 µL of medium were discarded and 10^4^ MOLT4–CCR5 cells were added to each well as target cells. At day 5, 100 µL of medium was replaced with fresh media. At day 14, the supernatant of each well was tested for p24 production using ELISA as previously described (35).

### VOA Sequence Amplification

Extraction of RNA and generation of cDNA and amplification of full-length *env* was performed as previously described (see extended data table 6 for a list of primers) (34, 36). The 1% 96-well E-Gels (Invitrogen) were used to visualize amplified *env* PCR products and select the bands with the expected HIV-1 envelope size. Selected PCR products were subjected to library preparation using Illumina Nextera DNA Sample Preparation Kit (Illumina) as previously described (34). Briefly, DNA was diluted in nuclease-free water to 10–20 ng per well and subjected to tagmentation. The Illumina Nextera Index Kit was then used to ligate tagmented DNA to barcoded sequencing adapters. Subsequently AmPure Beads XP (Agencourt) were used to purify DNA. Each library consisted of 96 different samples, which were subjected to paired-end sequencing using Illumina MiSeq Nano 300 (Illumina) cycle kits at a final concentration of 12 pM.

### Single-genome Amplification of Plasma Rebound Virus

Single-genome amplification (SGA) and sequencing of HIV-1 *env* genes was performed as previously described (37–39).

### Near Full-length Genome Amplification

CD4^+^ T cells were isolated from cryopreserved PBMCs and lymph node mononuclear cells (LNMC) using magnetic beads (Miltenyi). For genomic DNA extraction, we used Gentra Puregene Cell Kit (Qiagen), according to the manufacturer’s instructions. Near full-length sequencing (NFL) was done by an initial limiting-dilution semi-nested PCR amplifying the *gag* gene using the primers 3GagIN, 5GAGIN, 3GAGININ. If these primers failed, we used the primers GAGB5out, GAGB3out, GAGB5in, GAGB3in as previously described (8, 9) (extended data table 6). *Gag* PCR products were visualized using 1% 96-well E-Gels (Invitrogen). Dilutions with < 30% positive of the PCR wells were selected for further analysis. According to Poisson statistics, this dilution has over 90% probability of containing one HIV-1 DNA molecule in each PCR reaction. The NFL HIV-1 genome was amplified as a nested PCR with primers and cycling conditions as previously described (8, 9, 36). Briefly, the outer 9,064 bp PCR was performed using the primers BLOuterF and BLOuterR (extended data table 6) and High Fidelity Platinum Taq Polymerase (Invitrogen). For the nested PCR, 0.75 µl was transferred and the *env* gene amplified using the primers envB5out and envB3out. Wells containing an intact *env* gene were selected using 1% 96-well E-Gels (Invitrogen) and the corresponding outer PCR products were collected for further analyzes. NFL outer PCR products were subjected to a nested PCR to generate four segments A, B, C, D comprising overlapping parts of the genome. The PCR products were visualized on a 0.8% agarose gel to determine amplicon size. PCR products with the accepted size (A: 4,449 bp; B: 5,793 bp; C: 6,385 bp; D: 4,778 bp) were combined as either A+C, A+D, B+C or B+D and subsequently subjected to library preparation and sequencing using Illumina MiSeq Nano 300 (Illumina) cycle kits at a final concentration of 12 pM, as described above. Assembly and analysis of HIV-1 genome sequencing was performed as previously described (10).

### Identification of intact proviruses and construction of phylogenetic trees

To identify intact NFL sequences, we aligned assembled sequences to HXB2. Hereby we could identify premature stop codons, out-of-frame insertions or deletions (indel), or packaging signal (Ψ) deletions and mutations using custom Python scripts. Intact genomes were identified as sequences containing productive genes and the major splice donor (MSD) site. Sequences which had a deleted or mutated MSD site were categorized as Ψ-MSD deletion/mutation. Los Alamos HIV Sequence Database Hypermut tool was used to determine presence of APOBEC-induced G-A hypermutation in the remaining NFL sequences. Sequences which were not categorized as hypermutated, were considered defective due to indels/nonsense mutations. Trees combining all sequences for each individuals (intact and defectives) are shown in supplementary material (extended data figure 3). Maximum likelihood phylogenetic trees were constructed as previously described (10). To assess for cross-contamination of samples, we also generated a neighbor-joining (NJ) tree, which included all sequences obtained for the entire analyses and hereby confirmed that all sequences clustered correctly (extended data figure 4).

### Data availability

Sequence data generated in this study have been deposited in GenBank. Maximum likelihood phylogenetic trees showing all sequence names are in the supplementary material (extended data figure 5)

### Recombination analysis of *env* sequences

Multiple alignment of nucleotide sequences and the recombination analysis was performed as previously described (10, 20). Briefly, *env* sequences from NFL, VOA and SGA rebound sequences were analyzed for occurrence of recombination by the 3SEQ recombination algorithm (http://mol.ax/software/3seq/). Sequences with statistical evidence of recombination (i.e. rejection of the null hypothesis of clonal evolution) are represented in a circos plot (http://circos.ca/).

## Acknowledgements

We thank all study participants who devoted their time for the research and kindly donated their biological material. We thank Michel B. Hellfritzsch and Peter Ahlburg for assistance to obtain lymph node samples; Lene Svinth Jønke, Lilian Nogueira and Pilar Mendoza for laboratory assistance; Members of the Nussenzweig lab for productive discussions. L.K.V. is supported by a stipend from Aarhus University, Denmark. Y.Z.C. is supported by the National Institute of Allergy and Infectious Diseases of the National Institutes of Health under award number K23AI136578. O.S.S. is supported by a grant (#6110-00163) from the Danish Council for Independent Research. This work was supported by The Bill and Melinda Gates Foundation Collaboration for AIDS Vaccine Discovery (CAVD) grants OPP1092074, OPP1124068 (M.C.N.); the NIH grants 1UM1-AI100663 and R01AI-129795 (M.C.N.); the Einstein-Rockefeller-CUNY Center for AIDS Research (1P30AI124414-01A1); BEAT-HIV Delaney grant UM1 AI126620 (M.C.N.); and the Robertson fund. M.C.N. is a Howard Hughes Medical Institute Investigator. The funders had no role in study design, data collection and interpretation, or the decision to submit the work for publication

Author contributions: L.K. Vibholm, J.C.C. Lorenzi, Y.Z., Cohen, C-L. Lu, O. Søgaard and M.C. Nussenzweig designed the research. L.K. Vibholm, J.C.C. Lorenzi, T. Damsgaard, P.W. Denton, G.R. Teran, P.M. Estrada and Y. Ablanedo-Terrazas performed the research. L.K. Vibholm, J.C.C. Lorenzi, J.A. Pai, T.Y. Oliveira, J.P. Barton, M.G. Noceda, M. Tolstrup, O. Søgaard Y.Z. Cohen and M.C. Nussenzweig analyzed the data. L.K. Vibholm, Y.Z. Cohen, J.C.C. Lorenzi, O. Søgaard, P.W. Denton, M. Tolstrup and M.C. Nussenzweig wrote the manuscript.

The authors of this study declare no conflict of interest.

